# Conceptual structure and current trends in Artificial Intelligence, Machine Learning, and Deep Learning research in sports: A bibliometric review

**DOI:** 10.1101/2022.11.09.515813

**Authors:** Carlo Dindorf, Eva Bartaguiz, Freya Gassmann, Michael Fröhlich

## Abstract

Artificial intelligence and its subcategories of machine learning and deep learning are gaining increasing importance and attention in the context of sports research. This has also meant that the number of corresponding publications has become complex and unmanageably large in human terms. In the current state of the research field, there is a lack of bibliometric analysis, which would prove useful for obtaining insights into the large amounts of available literature. Therefore, the present work aims to identify important research issues, elucidate the conceptual structure of the research field, and unpack the evolutionary trends and the direction of hot topics regarding key themes in the research field of artificial intelligence in sports. Using the Scopus database, 1215 documents (reviews and articles) were selected. Bibliometric analysis was performed using VOSviewer and bibliometrix R package. The main findings are as follows: (a) the literature and research interest concerning AI and its subcategories is growing exponentially; (b) the top 20 most cited works comprise 32.52% of the total citations; (c) the top 10 journals are responsible for 28.64% of all published documents; (d) strong collaborative relationships are present, as along with small, isolated collaboration networks of individual institutions; (e) the three most productive countries are China, the USA, and Germany; (f) different research themes can be characterized using author keywords with current trend topics, e.g., in the fields of biomechanics, injury prevention or prediction, new algorithms, and learning approaches. AI research activities in the fields of sports pedagogy, sports sociology, and sports economics seem to have played a subordinate role thus far. Overall, the findings of this study expand knowledge on the research situation as well as the development of research topics regarding the use of artificial intelligence in sports, and may guide researchers to identify currently relevant topics and gaps in the research.

## Introduction

In general, both artificial intelligence (AI)^1^ and the subcategories of machine learning (ML) and deep learning have experienced a boom in recent years. AI is generally understood to be a type of human intelligence that is executed by machines. Machine learning is an approach to achieving AI, describing computer systems that are able to learn from examples or experiences without being explicitly programmed. Deep learning has also emerged as an important technique for implementing ML [1].

AI has already established itself in many situations and is now indispensable. For example, AI has experienced great success in facial recognition on smartphones [2], as well as speech processing, synthesis, and translation (e.g., Google Translate, Alexa, Google Assistant [3]), and has long since become routine in everyday life. It has proven itself in numerous real-world applications, including facial recognition [4], speech recognition [5], object recognition [6], malware detection [7], and spam filtering [8]. In addition, the main areas of application can be found in safety-critical fields of application in which errors have a tremendous impact (e.g., autonomous driving [9] or healthcare and medicine [10–13]).

The AI boom in recent years, along with the progressive development of the associated approaches, has been driven by various factors—including the availability (i.e., quality and quantity) of data, which decisively influences the success of AI models [14]. There have often been limited data in the past, but today, modern technologies often generate vast amounts of large and complex datasets [15]. In contrast to the original problem with the availability of data, there are now difficulties in coping with the sheer volume of data [16]. Parallel to the increase in the amount of available data, immense developments have taken place in the field of modeling, which have contributed to models becoming more and more accurate [1]. In addition, the implementation of ML models is becoming easier and more accessible for a large number of people thanks to user-friendly open-source programming libraries such as scikit-learn [17] or PyTorch [18]. Furthermore, hardware developments, the associated increase in available computing power, and the consequent acceleration of calculations play a decisive role [19]. These drivers have ensured that AI has opened up many new possibilities and gradually found its way into sports contexts and applications [20, 21]. Hence, the present work is based on an understanding of the field of AI in sports as the application of methods from the general field of AI in the context of sports science and practical sports issues.

The USA is considered to be a pioneer in using AI in sports, both in research and in practice. AI has been used for game and player analysis for several years, especially in American football and baseball [22]. For example, studies have shown that the analysis of various types of data can help support coaches’ tactical decision-making [23], as well as training and competition planning [24]. Furthermore, this kind of analysis has great potential for preventing injuries [20] which, in turn, is accompanied by a reduction in negative consequential costs and cancellations of competitions. Increasing numbers of major football clubs are also using AI and data analytics. First and foremost, FC Barcelona and TSG Hoffenheim have their own research labs (Barça Innovation Hub and TSG Research Lab, respectively). Liverpool are also implementing data analytics solutions and AI to support managerial decisions [25]. Even fan interactions can be monitored and analyzed for better performances [26]. On the topic of available data for analysis purposes, players are monitored via numerous different sensors during matches, competitions, and training. For example, with regard to the sub-disciplines of sports science, applications can be found in the fields of training science [24], biomechanics [27], and sports medicine [28]. Against the backdrop of the ever-increasing amounts of data, AI has become a key technology.

The research situation with respect to AI technologies in sports has also led to a large bibliographic dataset, which is impractical for manual investigation. There is a need to obtain optimal insights into the available publication data using appropriate analysis techniques, since it is crucial to analyze the relevant research and structures in depth to uncover existing knowledge, challenges, and research deficits in order to derive and justify new research necessities. A promising approach, which is becoming increasingly important in many fields of research, is bibliometric analysis. For example, bibliometric analyses are used for uncovering emerging trends, collaboration characteristics, and research constituents, as well as for exploring the intellectual structures of specific domains based on their respective literature base [29, 30]. Bibliometric techniques (as well as meta-analyses) seem especially promising and suitable for making sense of the large volumes of literature, as classic review methods need to focus on a manageable amount of literature [31]. Such techniques are superior to meta-analysis if the publications in question are very heterogeneous. Furthermore, unlike classic systematic literature reviews, which tend to be affected by the interpretation bias of scholars, bibliometric analysis (or meta-analysis) relies on more quantitative methods; thus, it can avoid or at least reduce such bias [29].

Regarding the basic review situation, several works have been published to date in the context of AI and sport-specific fields of its application [21,32–35]. In addition to AI topics, numerous general works can be found in the context of sports using bibliometric methods (e.g., football [36–38]; sports tourism [39, 40]; sports sustainability [41]; sports management [42]; sports nutrition [43]; sports science in general [44]). However, looking at different literature analysis methods applied in the context of sports-related AI research shows that only a few works have been published to date using bibliometric techniques. The authors of [45] analyzed literature regarding big data and artificial intelligence in sports using a bibliometric approach; however, their focus was restricted to articles in the CNKI database. The authors of [46] analyzed trends of AI applications in sports, with a focus on patents. Moreover, some works have used bibliometric techniques in the context of sports, focusing on topics that overlap with AI. For example, [47] analyzed publicly available baseball data in the context of big data characteristics. Furthermore, [48] investigated applications of wearable devices in sports using bibliometric techniques.

To the best of our knowledge, there is still a research deficit in the bibliometric analysis of available literature in the field of AI in sports using a broad literature base. Thus, it can be concluded that the overall scientific landscape has not been thoroughly analyzed thus far, and great potential remains.

Therefore, the present study aims to identify important research issues, reveal the conceptual structure, and unpack evolutionary trends and the directions of hot topics regarding key themes in the research field of AI in sports. Due to the presence of large bibliographic datasets, as well as the broad and heterogeneous literature in the context of machine learning research in sports, the application of a bibliometric analysis appears to be excellently suited to the intended object of investigation.

The following contributions of research constituents were analyzed using bibliometric methods through general performance analysis, and their relationships were identified using science mapping. Longitudinal development and current trends were further analyzed by looking at the occurrence of keywords over time and in the latest research publications. For in-depth background information on the applied analysis techniques, see [29], for example.

## Methods

For this study, data from the Scopus database were used (accessed on 6 September 2022). To map the most prominent subcategories of artificial intelligence, the terms “machine learning” and “deep learning” were included in the search, and “artificial intelligence” was the general term. It should be noted that the Scopus search engine is not case-sensitive and considers hyphens as spaces.

The document types selected were articles and reviews, as commonly performed in literature searches [40, 49], with a focus on works of the highest quality. As usual in bibliometric analyses, we focused on one language only (i.e., English). Due to the broad range of different subject areas addressed by publications in the context of sports, the exclusion of inappropriate literature based on subject areas of the database was not suitable for the aims of this work. Inappropriate literature in this context can be defined as publications that use the term “sport*” in the sense of a conceivable application of their research but do not carry out the research in the direct context of sports. However, in the present work, the focus was on works that are closely related to sports. We dealt with this problem using a literature search focusing on titles and the keywords, searching only for publications that included the term “sport*” in their title or keywords. Overall, this procedure allowed us to ensure that sports were the direct focus of the research, rather than a side issue. However, there was also the potential for inclusion of papers whose association with sports is debatable, depending on the individual and cultural understanding of the nature of sport. The final search procedure is shown in detail in Figure 1, resulting in 1215 documents from 428 journals (1157 articles, 58 reviews).

**Fig 1.**
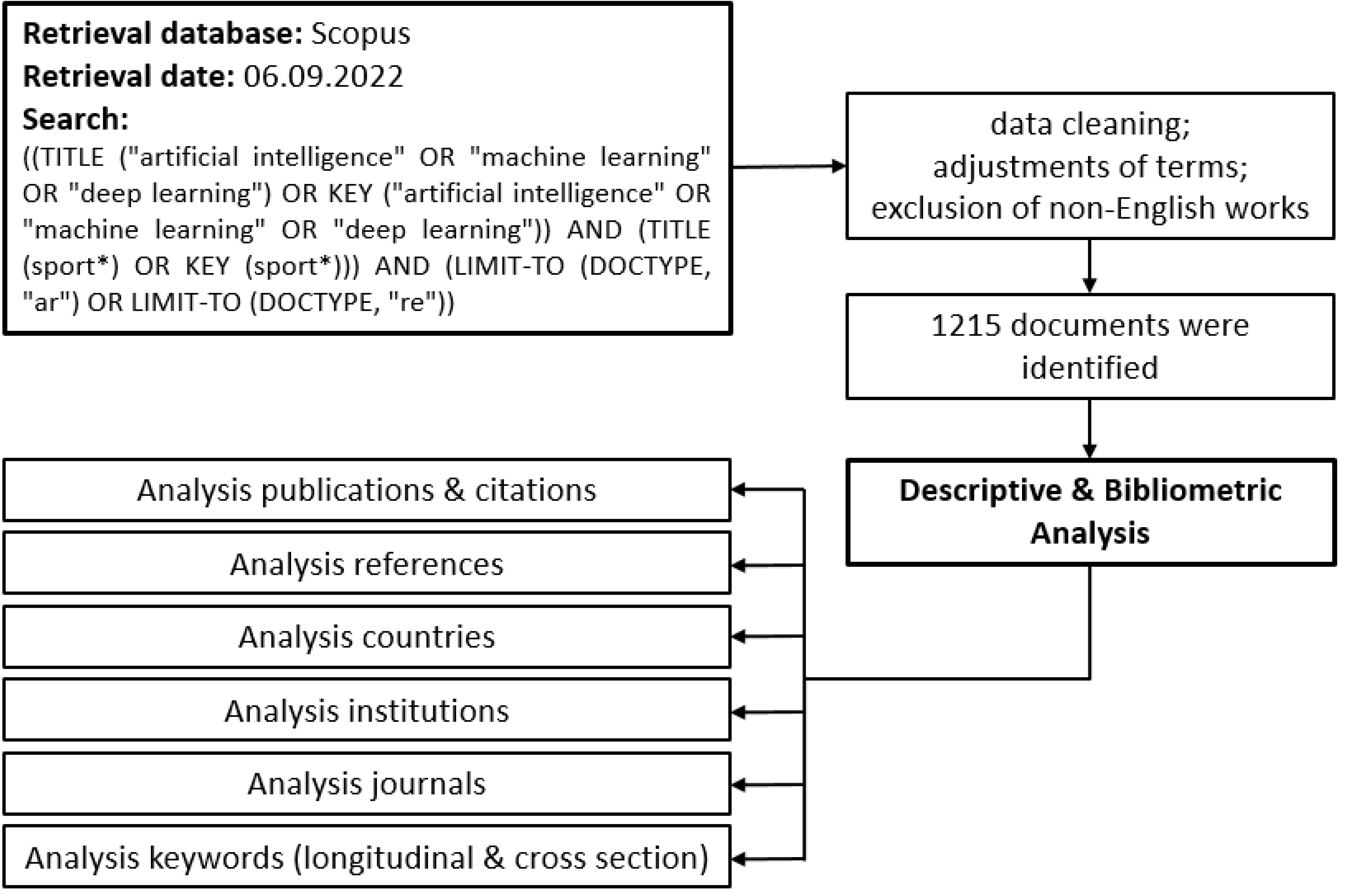
General workflow of the study.

Based on the final search, a csv file was exported. Erroneous or incomplete entries, as along with duplicates, were removed. Furthermore, abbreviations were replaced with their full designations (e.g., AI → artificial intelligence; ML → machine learning), while grammatical numbers were unified in the keywords (e.g., sport → sports) in the exported csv file. Incorrect and inappropriate keywords were manually removed.

Bibliometric analysis was performed using VOSviewer [50] and the R package bibliometrix, as well as the corresponding web interface biblioshiny [51]. VOSviewer and bibliometrix are commonly used in bibliometric research [49,52,53]. For a detailed overview of the software tools used, see [54]. Accordingly, the selection of the used software tools can be justified, since bibliometrix stands out due to the large number of different analysis methods available, while VOSviewer provides outstanding network visualizations.

In addition to the general performance analysis of the contributions of research constituents (e.g., articles, journals, institutions, authors, countries) to the considered field—using publication counts reflecting productivity and citations for measuring impact and influence [29]—the following science mapping methods were applied to identify the relationships between the research constituents: Co-authorship analysis focuses on revealing social interactions and relationships between authors, institutions (co-authorship institutions), and/or countries (co-authorship countries). Co-citation analysis identifies relationships among cited publications to reveal underlying themes in the research field. The co-occurrence of two publications in another publicatiońs reference list forms a connection between them [29]. As co-citation analysis focuses only on highly cited publications, bibliographic coupling was further applied to increase the visibility of recent and niche publications [29]. The analysis of bibliographic coupling shows which articles and authors are correlated with one another through multiple citations. A bibliographic coupling is present if two documents cite the same reference [55]. As it is recommended to use bibliographic coupling for specific timeframes [56], our analysis focused on the two recent periods 2019-2020 and 2021-2022.

Scopus uses the indexed keywords for the search; however, our keyword results are presented for the author keywords. This decision was based on our objective of starting from a broad, exploratory point of view in order to make the best use of any potentially relevant information and then focusing on aspects seen as relevant from the perspective of the scientific community. In addition to the general analysis of the total keyword occurrence, a longitudinal analysis of the keywords’ occurrence per year was also performed. This analysis was intended to reveal trending topics of AI research in sports based on the author keywords, which are generally sufficient to conclude topical aspects in a research field, as they are connected to the content of publication [57].

Furthermore, based on the author keywords, a thematic map of AI in sports was constructed. In general, the aim was to obtain insights into the current status of the field as well as to provide knowledge regarding the possible future development of research themes within the field. Therefore, the author keywords and their interconnections were clustered into themes. Characterization was performed through the properties density (i.e., measurement of the cohesiveness among the nodes, or internal cohesion; vertical axis) and centrality (i.e., measurement of correlation between different topics, or external cohesion; horizontal axis) [58]. The more relationships a node shows with others in the network, the greater its centrality and importance. The density representing the cohesiveness among the nodes gives clues as to the capability of a research field to develop and sustain itself [59].

## Results

### General performance analysis

In terms of the various Scopus subject areas, “Computer Science” (n = 741) ranked in first place, followed by “Engineering” (n = 517) and “Mathematics” (n = 242). The first publications were recorded in the year 1988 [60]. There was a steady increase in the annual number of publications, with the first major increase in 2012 and a second sharp and continuous increase from 2017 onwards (see Figure 2). Therefore, this research field can be thought of as being relatively young, especially in relation to the recent increase in publications. The context of the COVID-19 pandemic in the years 2020–2022 seems to have had little impact, with no visible drop in publications compared to other research fields, e.g., life sciences [61]. However, due in part to long publication times, it remains unclear whether the pandemićs effects will become apparent with a shift in the next few years.

**Fig 2.**
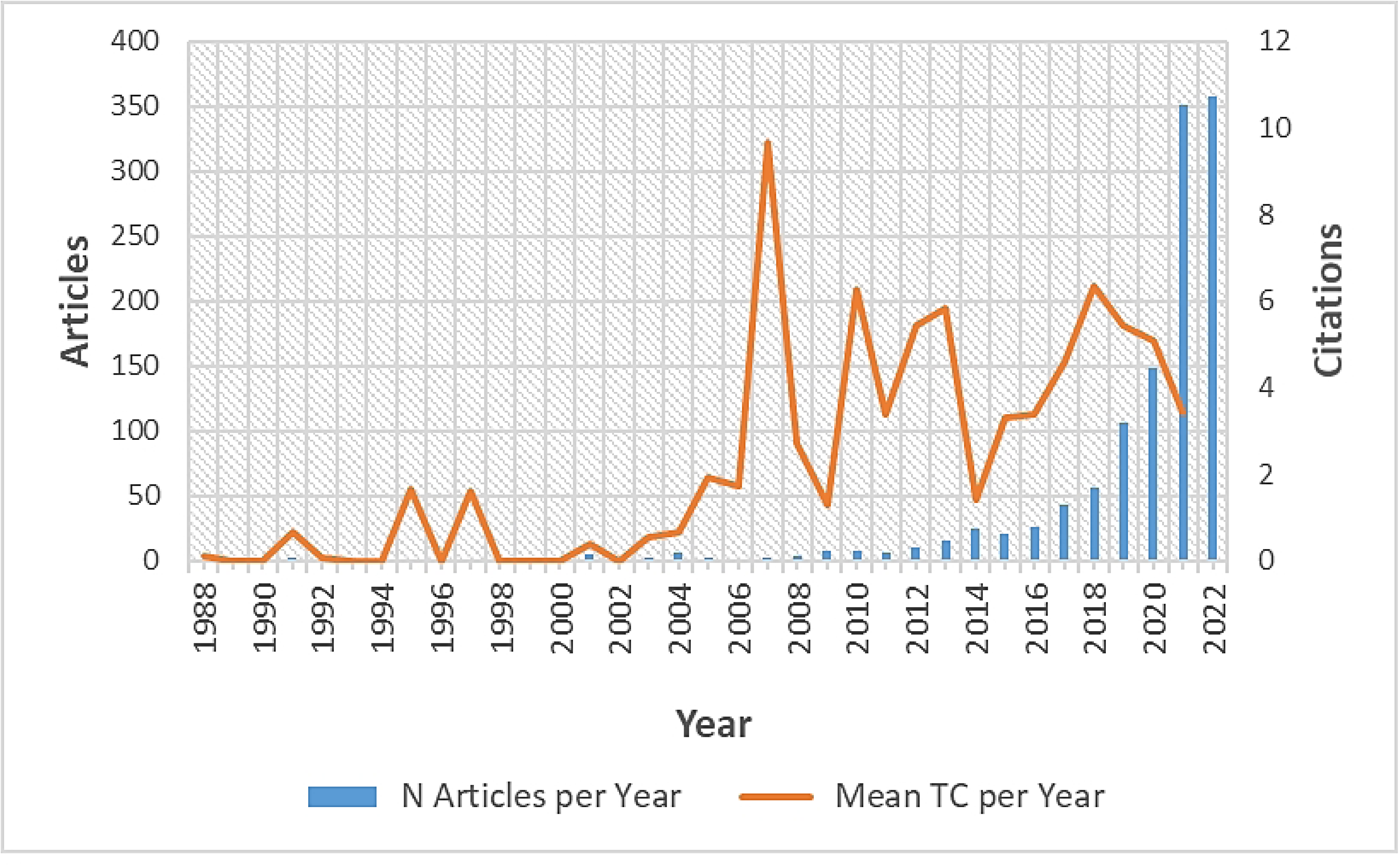
Evolution of the annual scientific production as the number of articles per year (left y-axis) and mean total citations per year (right y-axis). TC = total citations

Article citation analysis is the most widely used method to study the impact of authors, journals, and articles, as it identifies the key works in a research area [62]. Overall, on average, every document had 9.53 citations based on the global citation counts using Scopus data. Along with the total increase in the number of published documents over time, a parallel increase in the average number of citations per year was observable (see Figure 2).

An anomaly was detected for the year 2007 in terms of the total citations per document. This can be explained by the appearance of the most cited article “Checkers Is Solved” (Schaeffer et al., 2007) (see also: Table 1), which caused an atypical total citation count in relation to the early development of AI research in sports during that time. Checkers is a strategy board game that can be understood as a sport, similar to chess. In terms of citations per document, an unequal distribution was present, where the majority of works were only cited a few times. The proportion of papers with two or fewer citations was 53.33%. There are two potential reasons for this: the articles were published very recently, or they do not generate sufficient academic interest to be cited by the research community.

**Table 1.**
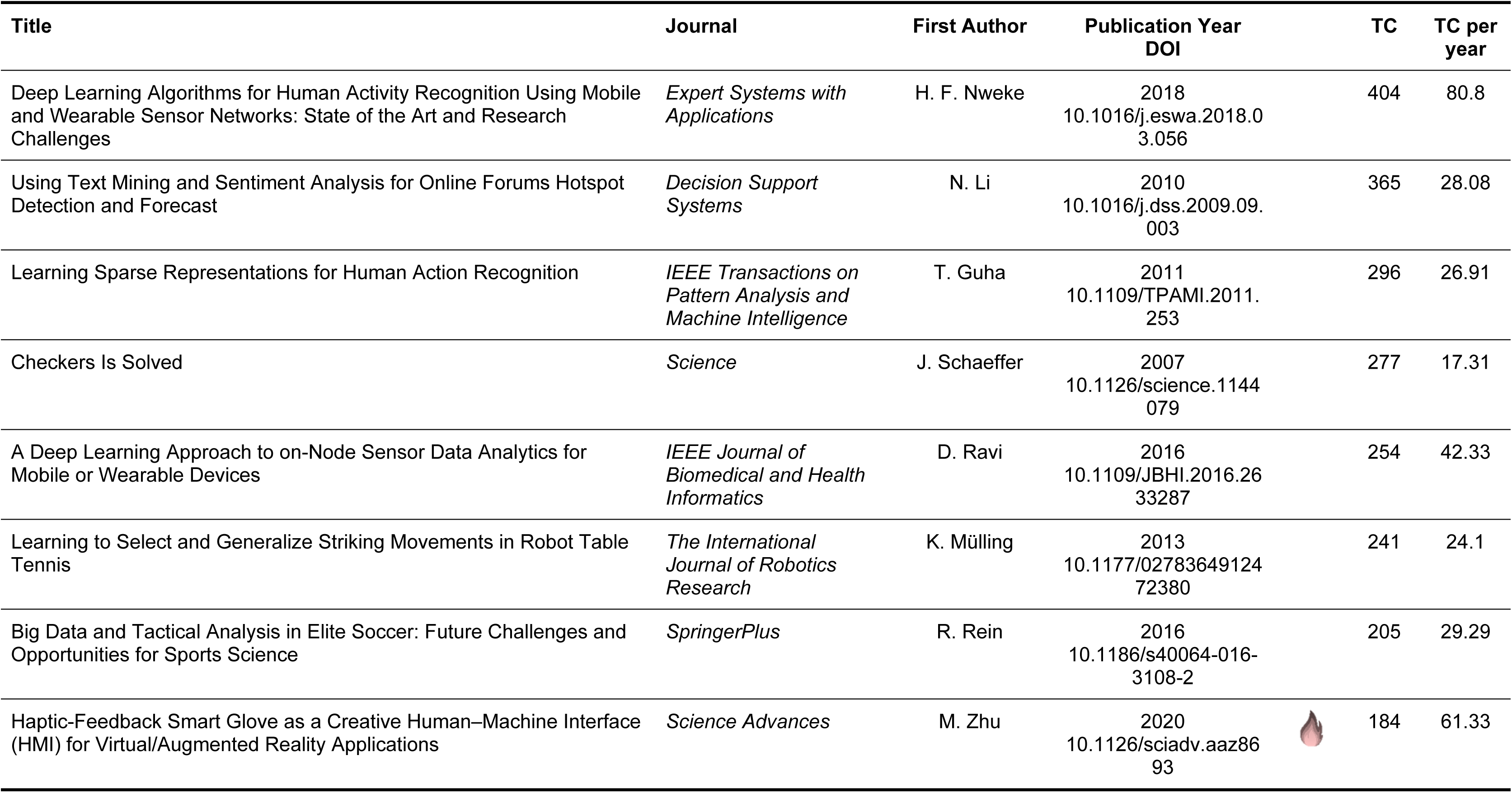
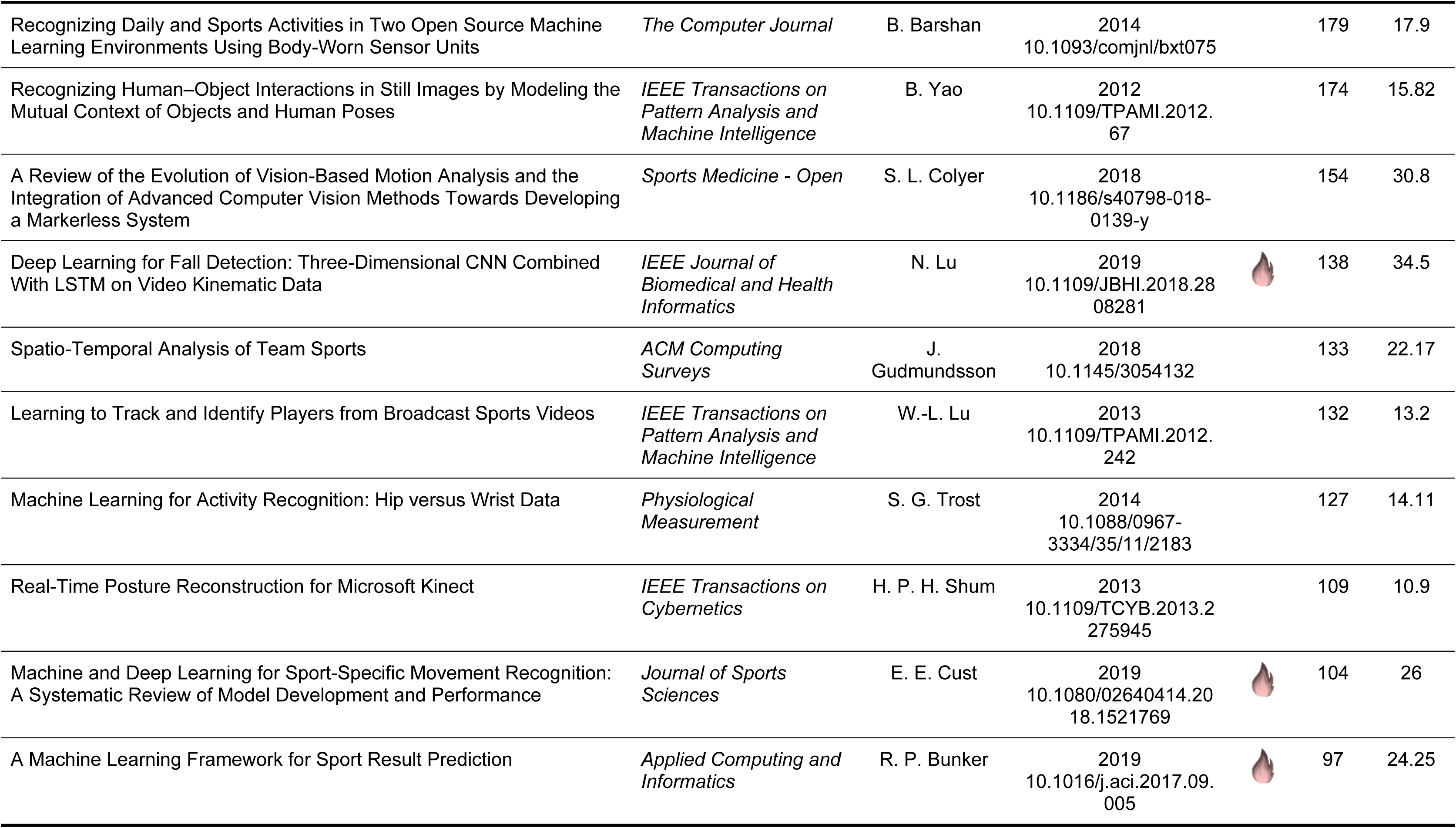
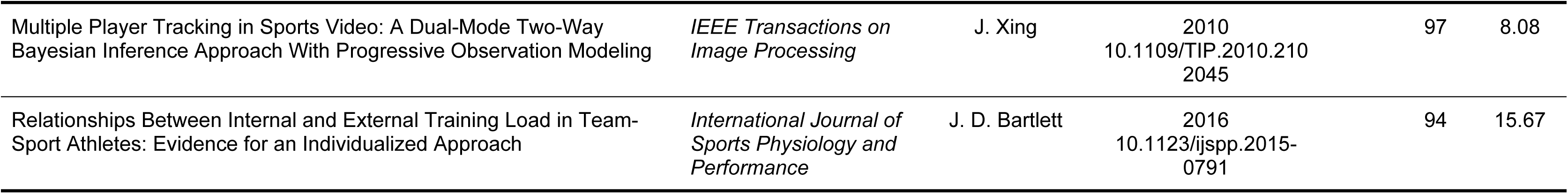
The top 20 most cited works. TC = total citations; normal. = normalized; 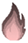 = “hot papers” published in the last four years.

Overall, the reviewed documents were published in 428 different journals. The 15 most productive journals in terms of the number of published documents are presented in Figure 3. The top 10 journals are responsible for 28.64% of all published documents. The most productive journal was *Computational Intelligence and Neuroscience* (h-index: 61; 2021 impact factor: 3.64), followed by *Sensors* (h-index: 196; 2021 impact factor: 4.35) and *IEEE Access* (h-index: 158; 2021 impact factor: 4.34). It is notable that the top-ranked journals include not only journals with specific aims and scopes, but also those with a broader possible range (e.g., *PLOS One*). However, journals that focus on information technology and engineering comprise the majority. This indicates that AI in sports is interdisciplinary, because most journals do not focus on sports. The *Journal of Sport Science* is only ranked 10th, which can be taken as a sign that AI is not the main topic in sports science at all.

**Fig 3.**
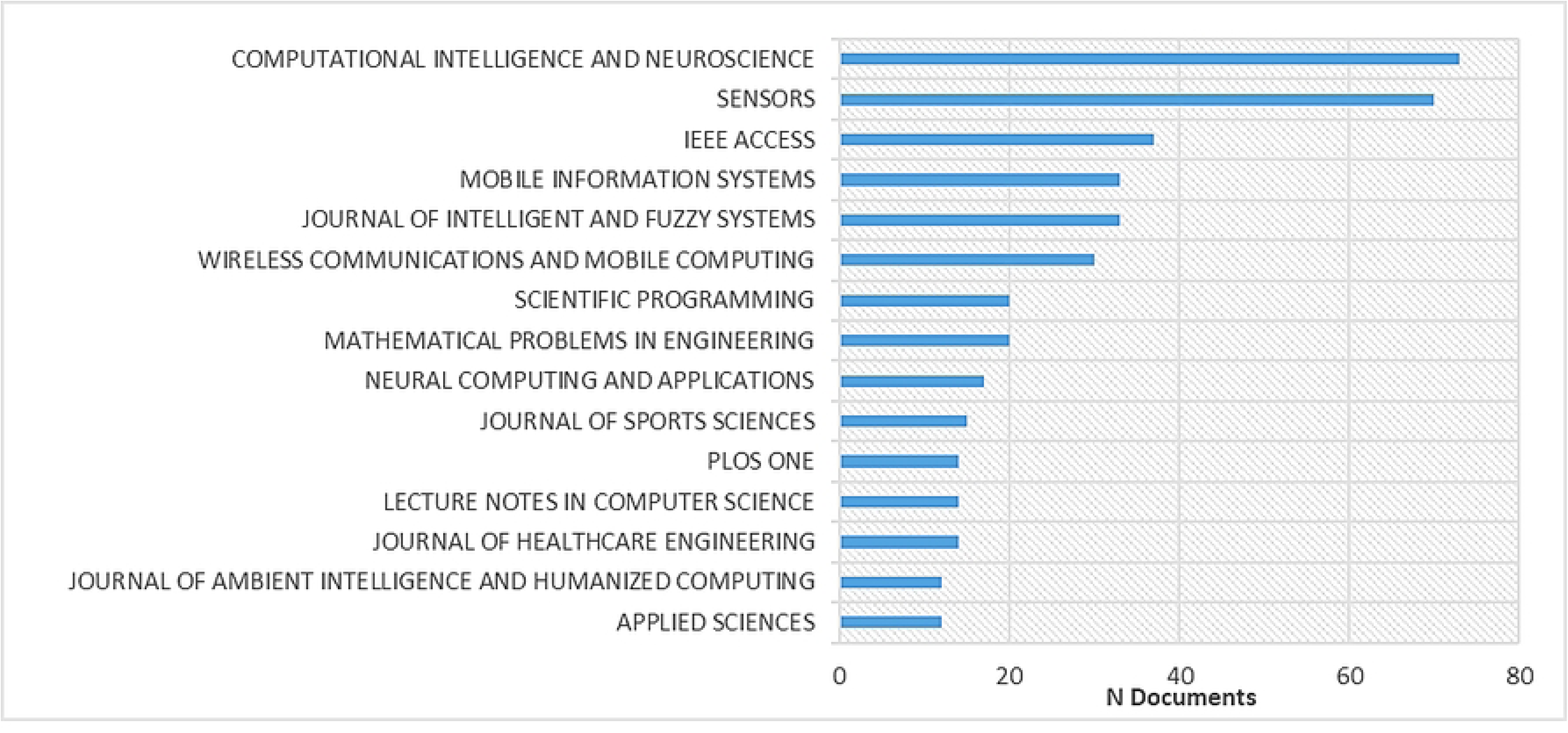
The top 15 most productive journals for research articles in the context of sports and artificial intelligence.

### Top 20 overall most cited works

The top 20 most cited works (see Table 1) account for 32.52% of the total citations. The most common theme in the top 20 most cited works is human activity or action recognition (e.g., [32,63–67]), followed by predictive systems (e.g., [68, 69]) and systems for object or player tracking (e.g., [70–72]). Only one study investigated AI in the direct context of motion analysis [73]. Most studies developed an algorithm or a new methodological approach (e.g., [74–76]). The other studies focused on reviewing the literature [32,63,70,73,77]. The focus of the data was mainly on wearables (e.g., [63,65,67,76]), followed by visual data (e.g., [71–73,78]). Team sports were analyzed more frequently compared to other sports (e.g., [69,70,77]).

The publications that were a maximum of four years old and featured among the top 20 most cited publications in this context (referred to here as “hot papers”; see also: Table 1) could take on special importance in future research if they continue to be cited over the next few years.

### Countries, organizations, and global structures

AI in sports is a global research topic. Analysis of the total authorship data of each document showed that 128 countries contribute to the research on this topic. However, China leads in terms of the total number of publications by a considerable margin, followed by the USA and Germany (see Table 2). On average, each publication has 3.65 co-authors. Regarding the mean numbers of authors from the most productive countries, China has the fewest authors per document. This can be explained by the fact that 27.33% of the publications with at least one Chinese author had only one author; in comparison to the USA and Germany—with 7.38% and 2.99% single authorship, respectively—this is a relatively high proportion. It should be noted that 21.15% international co-authorship is also present; therefore, a single publication can be counted in several countries.

**Table 2.** Top 10 most productive countries based on the authoŕs countries, total citations (TC) per country based on document citations, and mean number of authors per country based on the authorś countries.

| Countries | N publications | TC | Mean number of authors |
| --- | --- | --- | --- |
| China | 1244 | 1723 | 2.72 |
| USA | 591 | 1031 | 5.27 |
| Germany | 204 | 842 | 4.40 |
| UK | 170 | 671 | 4.39 |
| Australia | 169 | 815 | 4.54 |
| South Korea | 149 | 395 | 3.56 |
| Japan | 134 | 139 | 5.39 |
| Spain | 134 | 159 | 5.02 |
| Canada | 125 | 796 | 4.23 |
| India | 112 | 107 | 3.76 |

With 38 articles, the Hospital for Special Surgery (New York City, New York) takes first place among the most productive organizations, followed by Victoria University (Melbourne, Australia) and Ghent University (Ghent, Belgium) (Figure 4a). Surprisingly, although China is the most productive country in terms of publication output, the leading Chinese university only ranks in 6th place (Beijing Sport University). This indicates that many institutions in China seem to be involved in the publications without any individual institution being particularly active. The aforementioned high proportion of single authorship in Chinese publications, without collaborations between institutions, also appears to be related to this.

**Fig 4.**
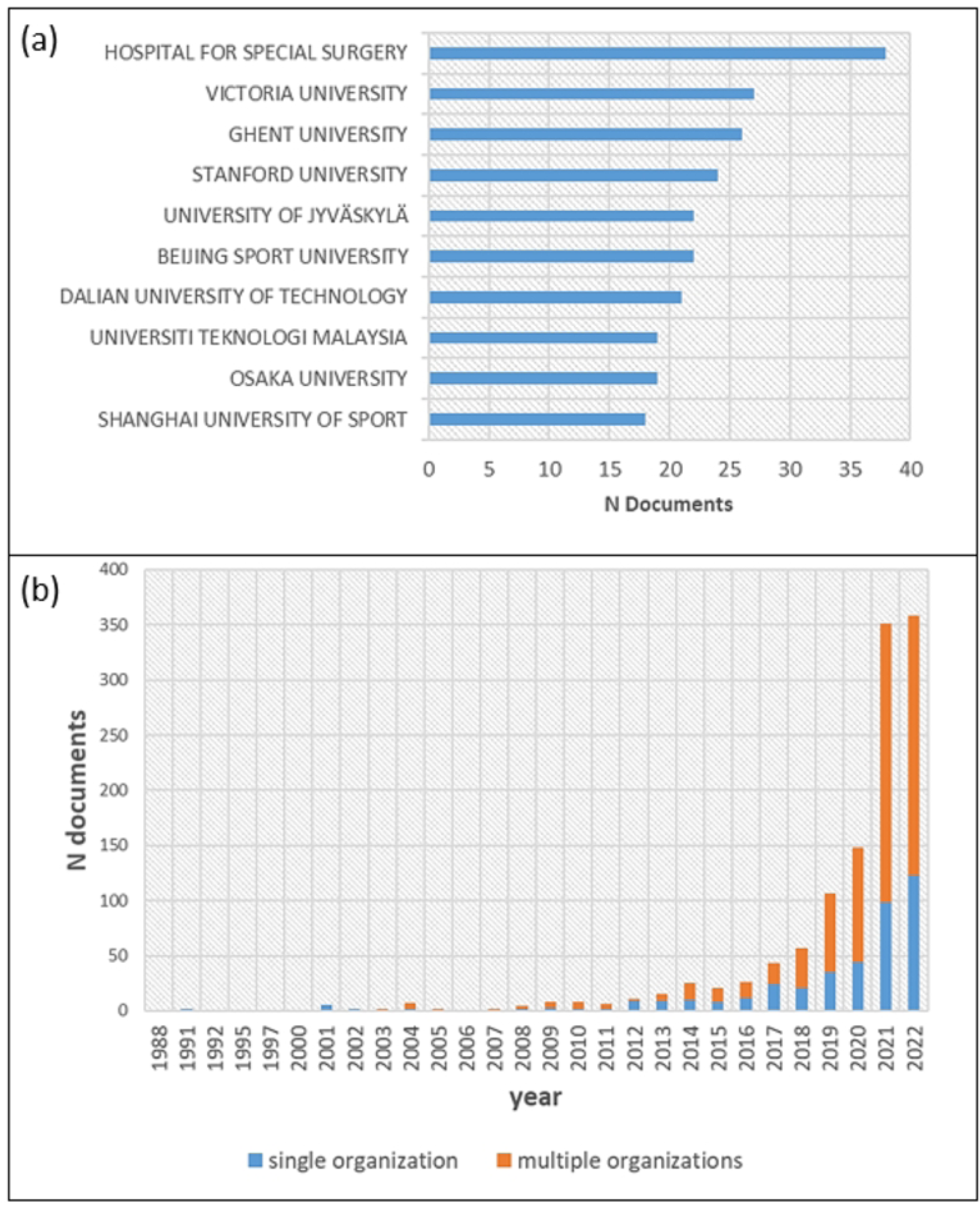
(a) Top 10 most productive organizations based on authors’ affiliations. (b) Distribution of publications according to the involvement of one vs. multiple universities.

It is notable that the trend toward multi-university publications has continued to increase, especially in recent years (Figure 4b). This could be interpreted as a sign that AI in sports has moved out of its initial niche status. An examination of the collaboration between organizations (Figure 5) shows that publications are often produced through cooperation between individual universities. There are larger associations of collaborating universities as well as islands formed through small collaboration networks. The resulting networks may be due to the need to bundle expertise, since individual universities mainly provide sporting or IT expertise, which must be brought together to process the research object in the sense of interdisciplinary or transdisciplinary research. According to the existing small, unconnected collaborative networks, networking still has great potential insofar as there is a fit in terms of the content and methodology of the research institutes.

**Fig 5.**
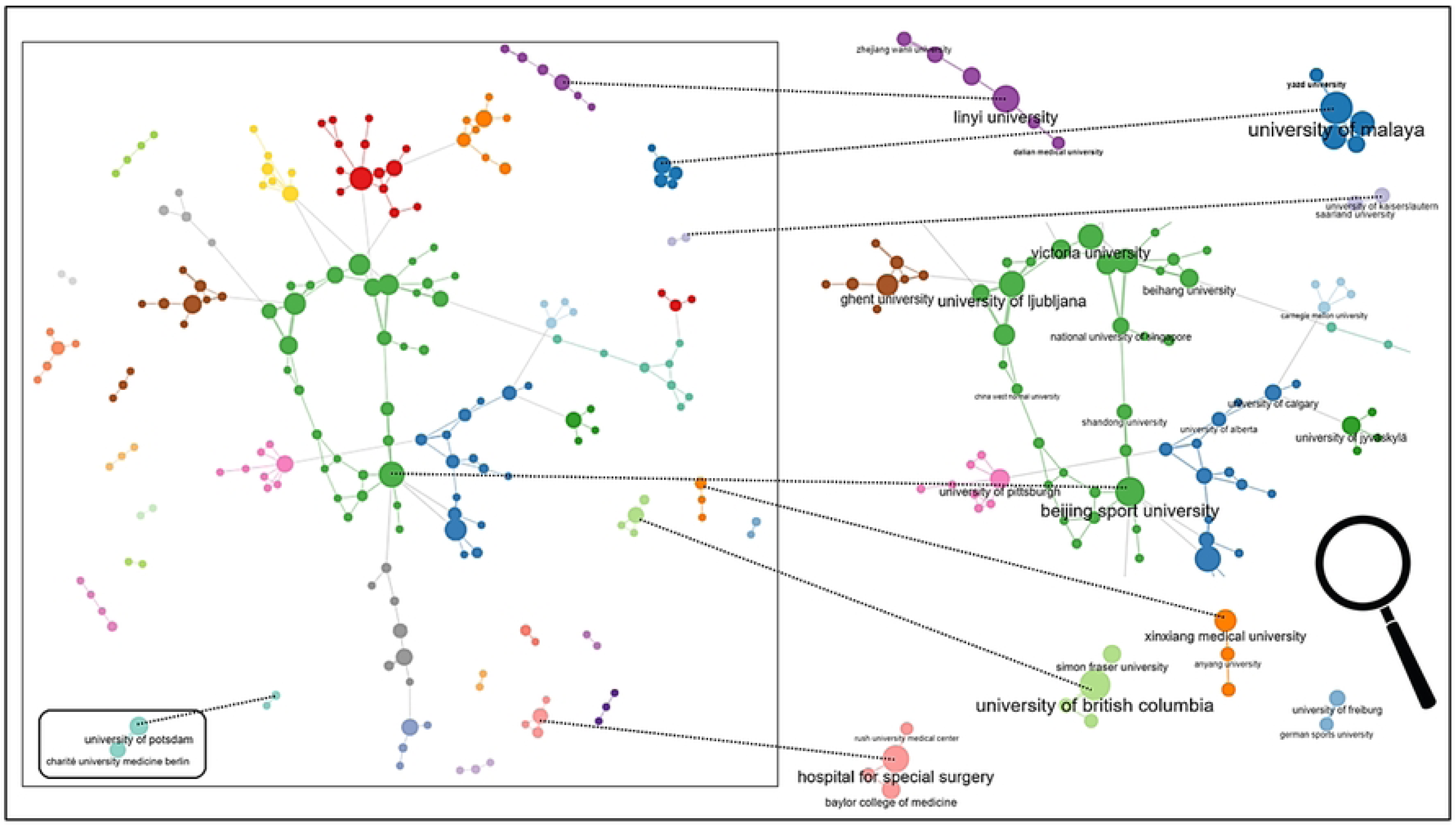
Collaboration network organizations using the bibliometrix package (parameters: Walktrap clustering algorithm, 500 nodes displayed). Exemplary subnetworks are also presented, zoomed in with the names of exemplary organizations for visualization reasons.

Examination of the international collaboration, which indirectly depicts the connections of the authors from different countries, shows that five clusters of collaboration groups are present (Figure 6). The top three most productive countries (i.e., China, the USA, and Germany) are in different clusters. In the blue cluster, the most productive countries are China, South Korea, and India. In the yellow cluster, the USA is the most active country, followed by Spain and Canada. In the red cluster, the most productive countries are Germany, Turkey, and Japan. In the green cluster, the United Kingdom is the leader, followed by Australia and Slovenia. It is striking that geographic proximity is a strong explanatory factor but is not necessarily related to cluster affiliation. For example, Germany collaborates strongly with the nearby countries of the Netherlands and Belgium, but also with Japan. The same applies to the cooperation between the USA and Spain.

**Fig 6.**
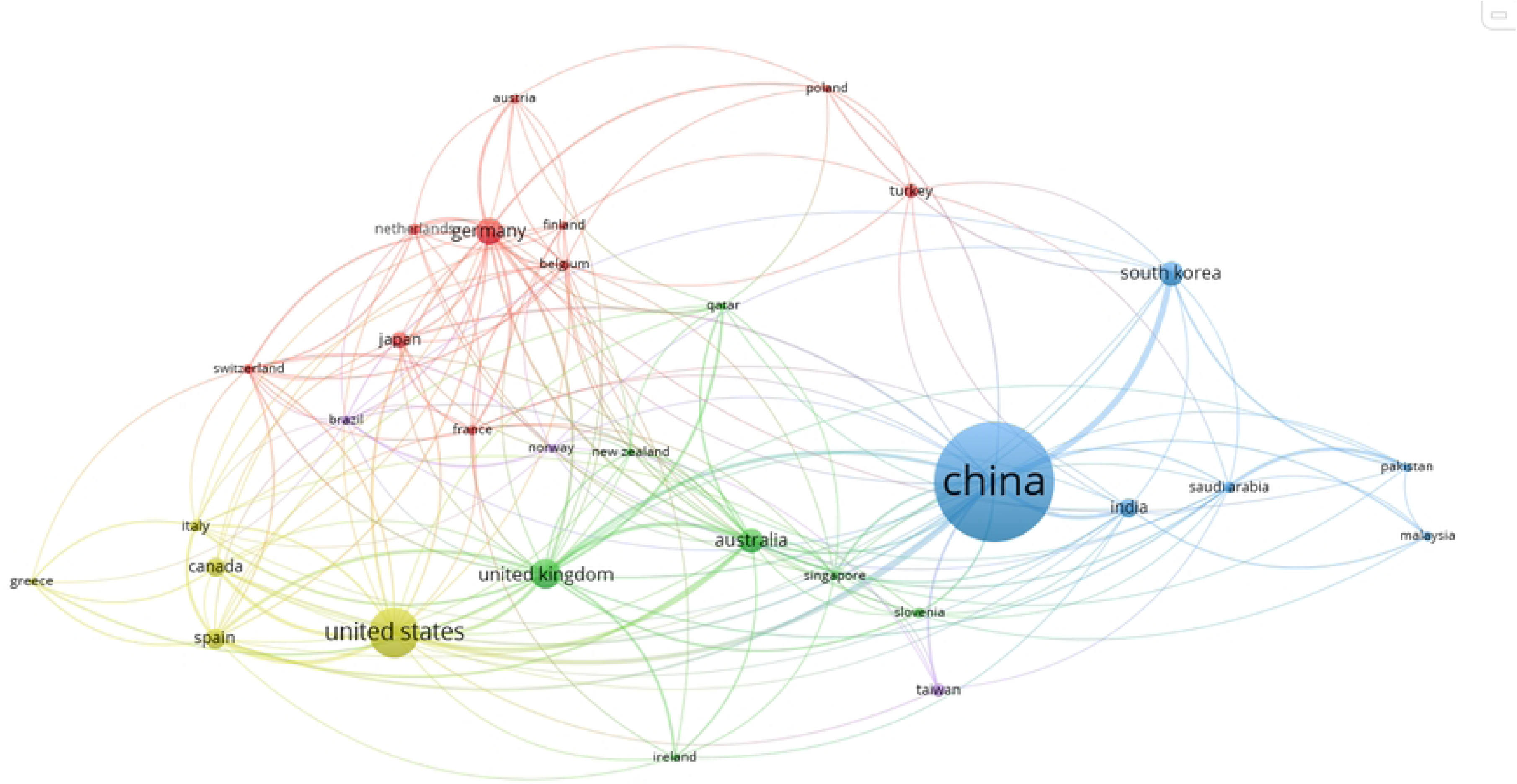
Collaboration world map/co-authorship countries: The minimum number of documents per country was set to 10, resulting in 31 countries. Cluster memberships are distinguished by color. The node size reflects the productivity of the country based on the number of publications in collaboration with other countries; the shorter the distance between the nodes, the more closely connected the countries are (based on co-authored articles).

Substantial collaboration is visible between the different clusters. However, the collaboration between China and the other main clusters is lower than the collaboration of the remaining clusters. Using the terms of the World Bank for the classification of the countries [79], it seems that high-income countries and middle-income countries collaborate more within each cluster.

### Authors and social structures

The research field of AI in sports is characterized by a large number of authors. In total, 3411 different authors published their works during the period of time investigated in this review. Only 188 authors of single-authored publications were present. On average, each publication shows 3.65 co-authors, and 21.15% of the papers are written by international author groups. The majority of the most productive authors are located in China. Due to the relatively high level of correspondence between names according to the country-specific characteristics in China [80], the most productive authors are not presented here, so as to rule out any possible errors or injustice, since authors were not identified on the basis of a clearly identifiable ID (e.g., ORCID ID) but on the basis of their first and last names which, due to the abovementioned situation, may have led to an incorrect merging of authors with the same first and last names. However, as it is reported that co-citation analysis is relatively immune to author name ambiguity issues [81], the corresponding mapping was performed to reveal the social structures of the authors (Figure 7).

**Fig 7.**
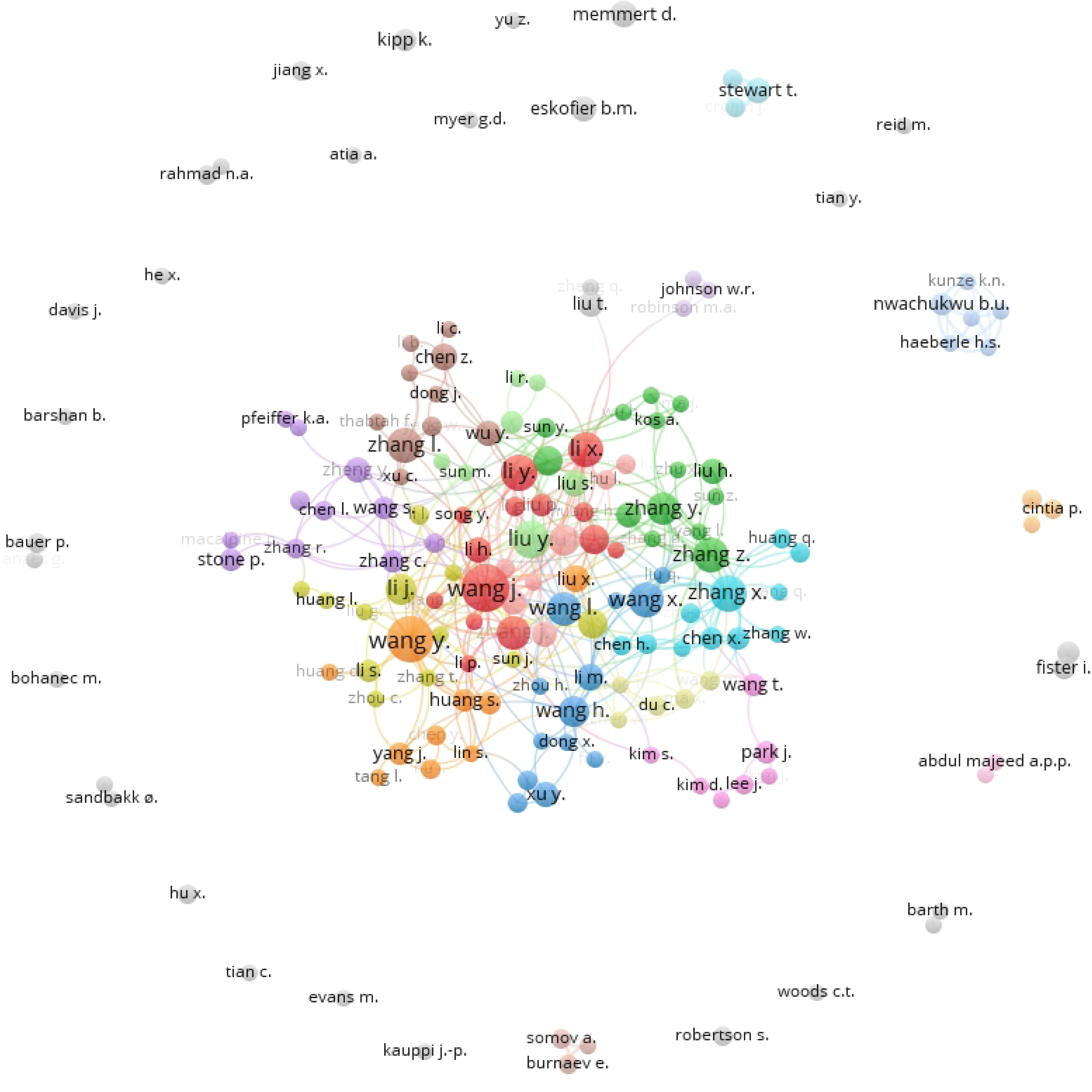
Co-authorship networks using VOSviewer: The minimum number of documents per author was set to 3, resulting in 172 authors. Cluster memberships are distinguished by color. The node size (author) reflects the number of received citations; the shorter the distance between the nodes, the more closely connected they are (based on co-citations).

In the center of Figure 7, we can see a large number of strongly networked and collaborating authors who come from China in particular. In the periphery, however, some authors do not show any networking, or small groups are formed that are not further connected to other authors. Unused potential in networks could be the reason for these islands’ formation. On the other hand, it also seems conceivable that the small, separate groups are pursuing very specific research areas and, therefore, there are no networking opportunities with researchers from other subject areas with methodological or content-related similarity.

### Co-citation analysis and Bibliographic coupling

The co-citation analysis revealed the presence of six clusters (see Figure 8). Two clusters were not connected with other clusters (shown in green and blue). The overall most cited publications in the clusters were those of Breiman (2001) with the random forest algorithm [82] and Hochreiter and Schmidhuber (1997) with long short-term memory (LSTM) [83].

**Fig 8.**
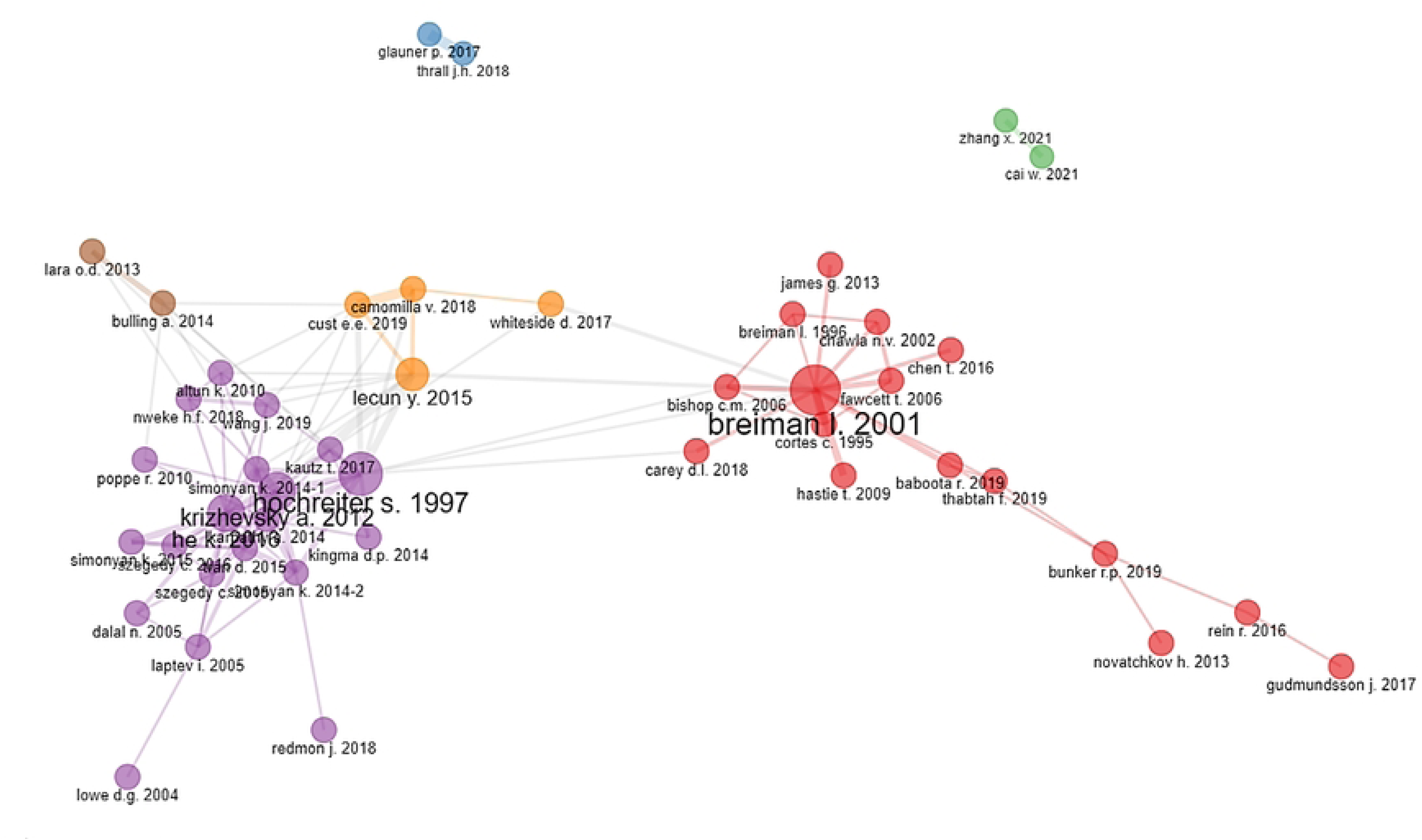
Co-citation analysis of documents using the bibliometrix R package: Leading eigenvalues were used as a clustering algorithm. Isolated nodes were removed. Fifty nodes are displayed. Cluster memberships are distinguished by color. The node size (article) reflects the number of received citations; the shorter the distance between the nodes, the more closely connected they are (based on co-citations).

Focusing on the main topics of each cluster, it is hard to find significant differences. The first cluster, shown in red, could be grouped under primary algorithmic and machine learning literature—most notably the paper of Breiman (2001) [82].

Many papers in the purple cluster deal with deep convolutional neural networks as well as human action or image recognition. Furthermore, the works of Zhang et al. (2021) [84] and Cai et al. (2021) [85] (green cluster), which are not connected to other clusters, can be attributed to image segmentation. The publications in the orange cluster (e.g., LeCun et al., 2015) [86] mainly deal with the themes of wearables and deep learning.

The results of the bibliographic coupling are presented in Figures 9 and 10. A look at the most frequently cited publications in the recent period (Figure 9) shows that only a few thematically distinguishable clusters can be formed for the years 2021 and 2022. With a minimum of three citations, the bibliometric analysis obtained 14 clusters. The first cluster, shown in red, contains 17 items, mainly concerning the themes of machine learning applications in the context of sports and health, as well as decision-making; in this cluster, the most representative paper was published by Cao et al. (2021) [87]. The green cluster includes 16 items and could be summarized under the topic of motion recognition; Cui et al. (2021) [88] published the most cited paper in this cluster. The blue cluster consists of 14 items; most of these publications address the topic of AI in medicine; the study of Ramkumar et al. (2021) [89] was cited the most, with 20 citations.

**Fig 9.**
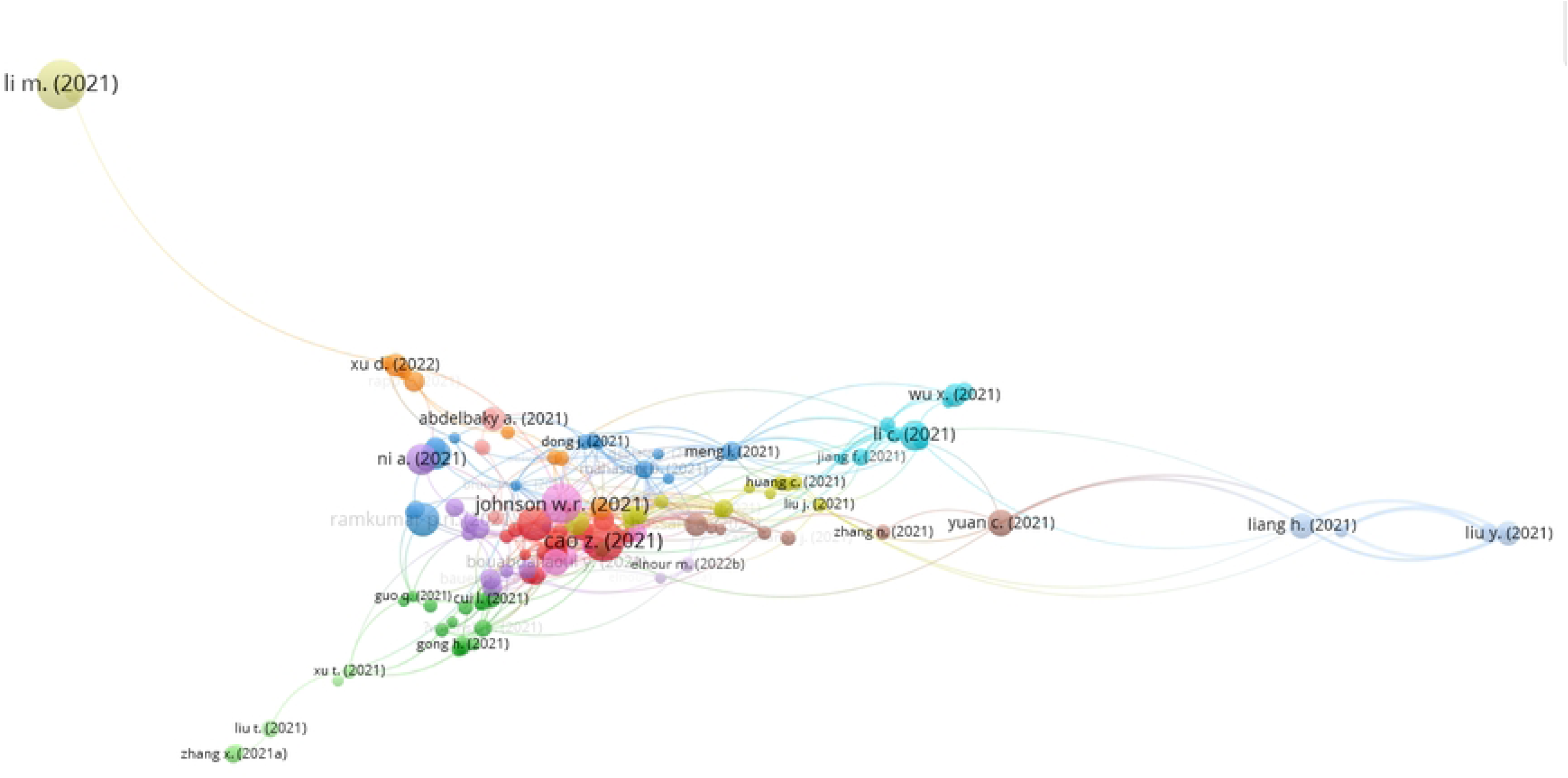
Bibliographic coupling for the years 2021 and 2022 using VOSviewer: The minimum number of citations per document was set to 3 (180 out of 714 selected). Only the largest set of connected items is displayed (129 items). Cluster memberships are distinguished by color. The node size (article) reflects the number of citations received; the shorter the distance between the nodes, the more closely connected they are (based on coupling).

**Fig 10.**
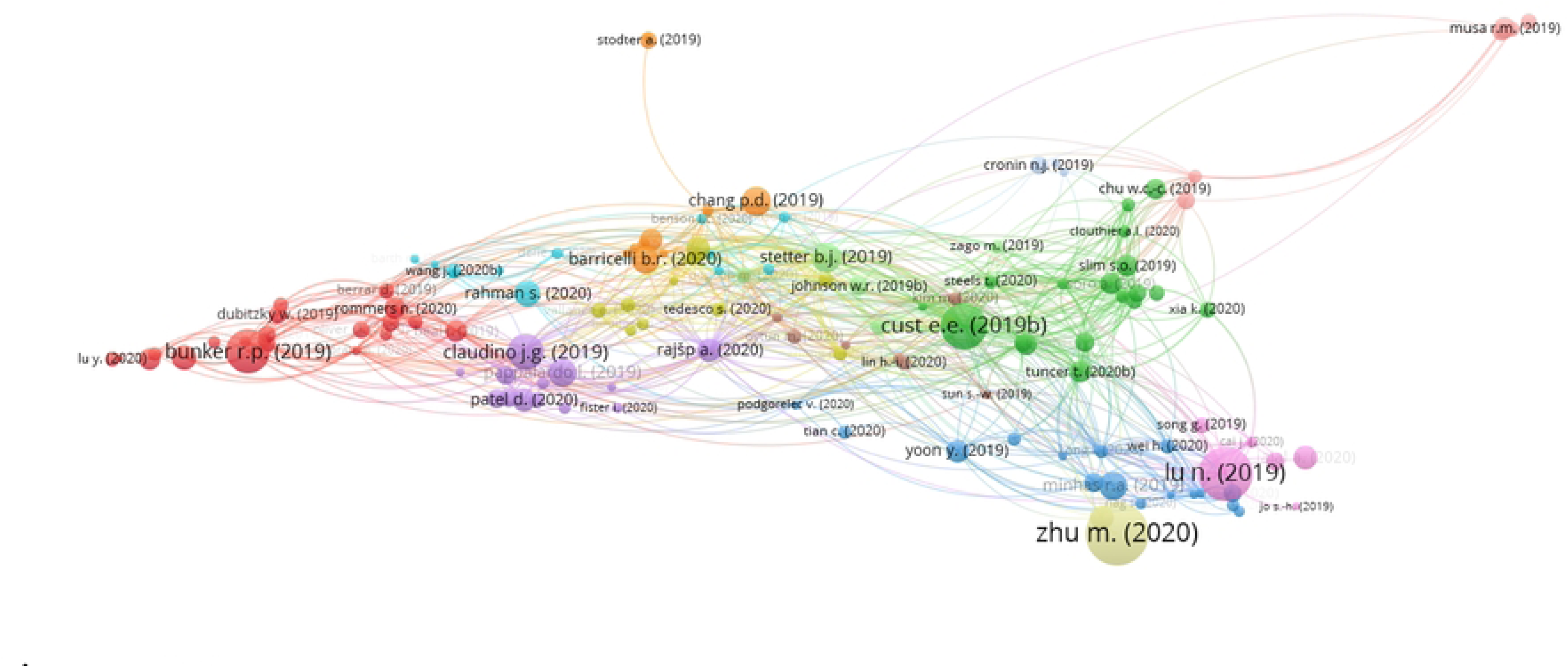
Bibliographic coupling for the years 2019 and 2020 using VOSviewer: The minimum number of citations per document was set to 5 (163 out of 257 selected). Only the largest set of connected items is displayed (135 items). Cluster memberships are distinguished by color. The node size (article) reflects the number of received citations; the shorter the distance between the nodes, the more closely connected they are (based on coupling).

Thirteen clusters were obtained via bibliometric analysis for 2019 and 2020 (see Figure 10). The red cluster contains 21 items, most of which can be assigned to the topic of team sports; the most cited paper in this cluster was published by Bunker et al. (2019) [68]. The cluster shown in green, with 16 items, can be summarized under the topic of sports recognition or recognition methods; the most representative paper in this cluster was published by Cust et al. (2019) [32]. The blue and purple clusters pertain to research with detections/tracking and big data, as well as machine learning; the most cited papers in the blue cluster were authored by Yoon et al. (2019) [90] and Minhas et al. (2019) [91]; in the purple cluster, the review by Claudino et al. (2019) [20] was cited the most.

### Keywords / Themes: general trends and longitudinal evolution

The overall occurrence of the 25 most frequently occurring author keywords is shown in Figure 11. The top four keywords map the search terms used for the Scopus search. It is notable that “machine learning” is the most frequently used term; the subarea “deep learning” is second, followed by “artificial intelligence”. A possible reason for this order is the more precise and specific scientific language [92] that specifically names the subfields instead of speaking of AI in a general way. The fact that the keyword “sports” only came in 4th place is a sign that the authors do not use sport as a keyword, in contrast to Scopus. This could be explained by two possible mechanisms: On the one hand, it is possible that authors do not use sport itself as a key term, referring instead to football, biomechanics, injury prevention, etc., as subsets of sport. Thus, sport(s) would be too general or too obvious as a keyword. On the other hand, it is also possible that authors do not see to the relevance of sport in their research compared to the AI topics.

**Fig 11.**
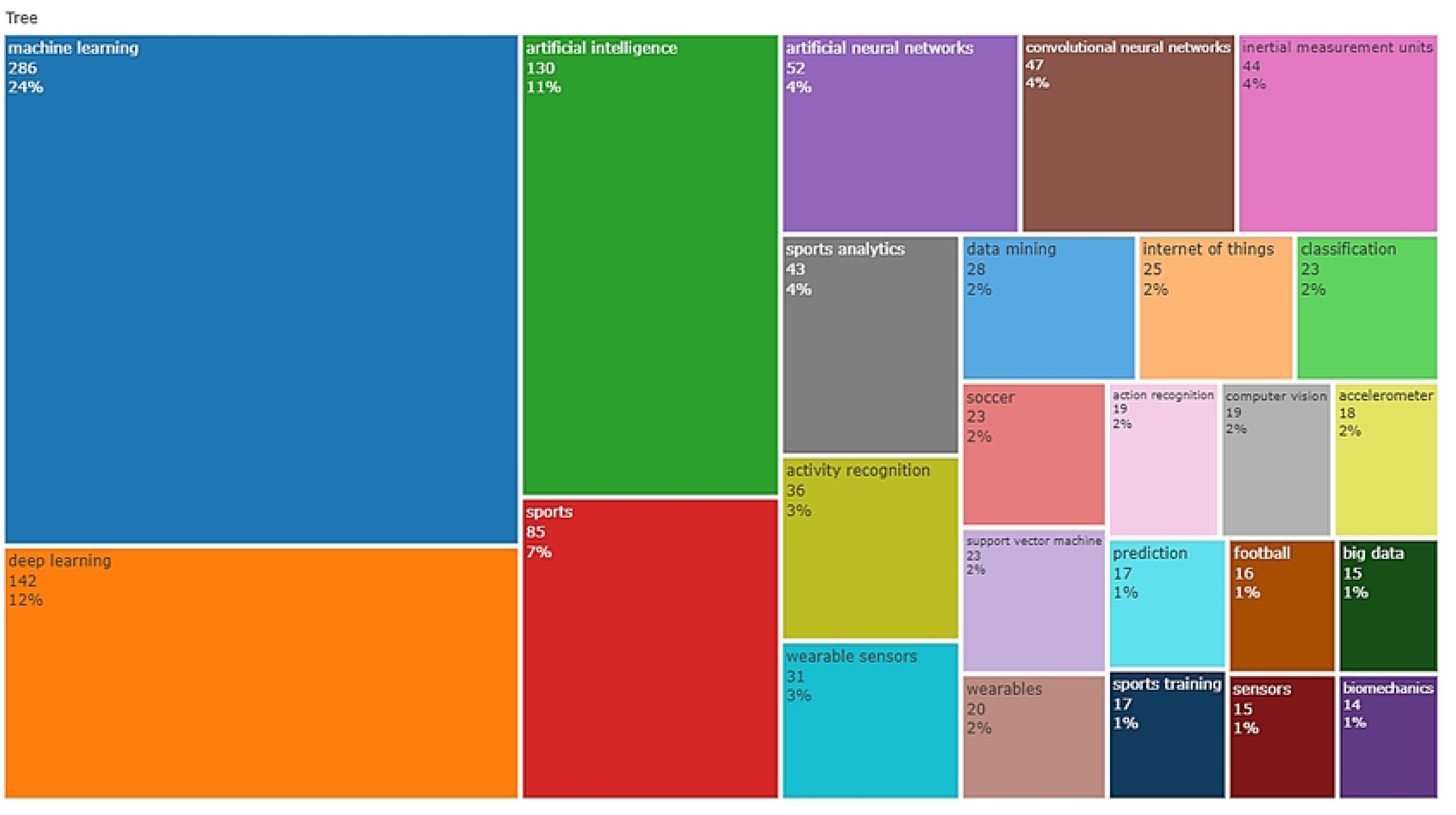
TreeMap of the 25 most frequently occurring author keywords

The keywords can be manually grouped according to their meaning. The keywords “artificial neural networks”, “convolutional neural networks”, and “support vector machine” can be subsumed under the term *algorithms* (a). The keywords “inertial measuring units”, “wearable sensors”, “accelerometer”, “wearables”, and “sensors” can be assigned to *measuring methods or tools* (b). It is striking that all of these methods can also be used outside of laboratory settings in the field and make huge amounts of data relatively easily accessible, which favors the emergence of “big data” characteristics [93]. Specific *AI tasks* (c) are mapped with the terms “classification”, “activity recognition”, “action recognition”, and “prediction”. Different *research fields* (d) are mapped with the terms “sports analytics”, “computer vision”, “data mining”, “internet of things”, and “biomechanics”. Of the 25 keywords shown, only “soccer” and “football” name specific *sports categories* (e). A closer analysis shows that football is used synonymously with soccer in most of the publications included, in accordance with country-specific conventions. Other sports were also mentioned as specific keywords; however, their frequency was significantly lower (see Figures 12 and 13). Thus, it can be seen that the sport of soccer/football is very dominant, which can also be explained by the significant financial interests behind this sport [94].

**Fig 12.**
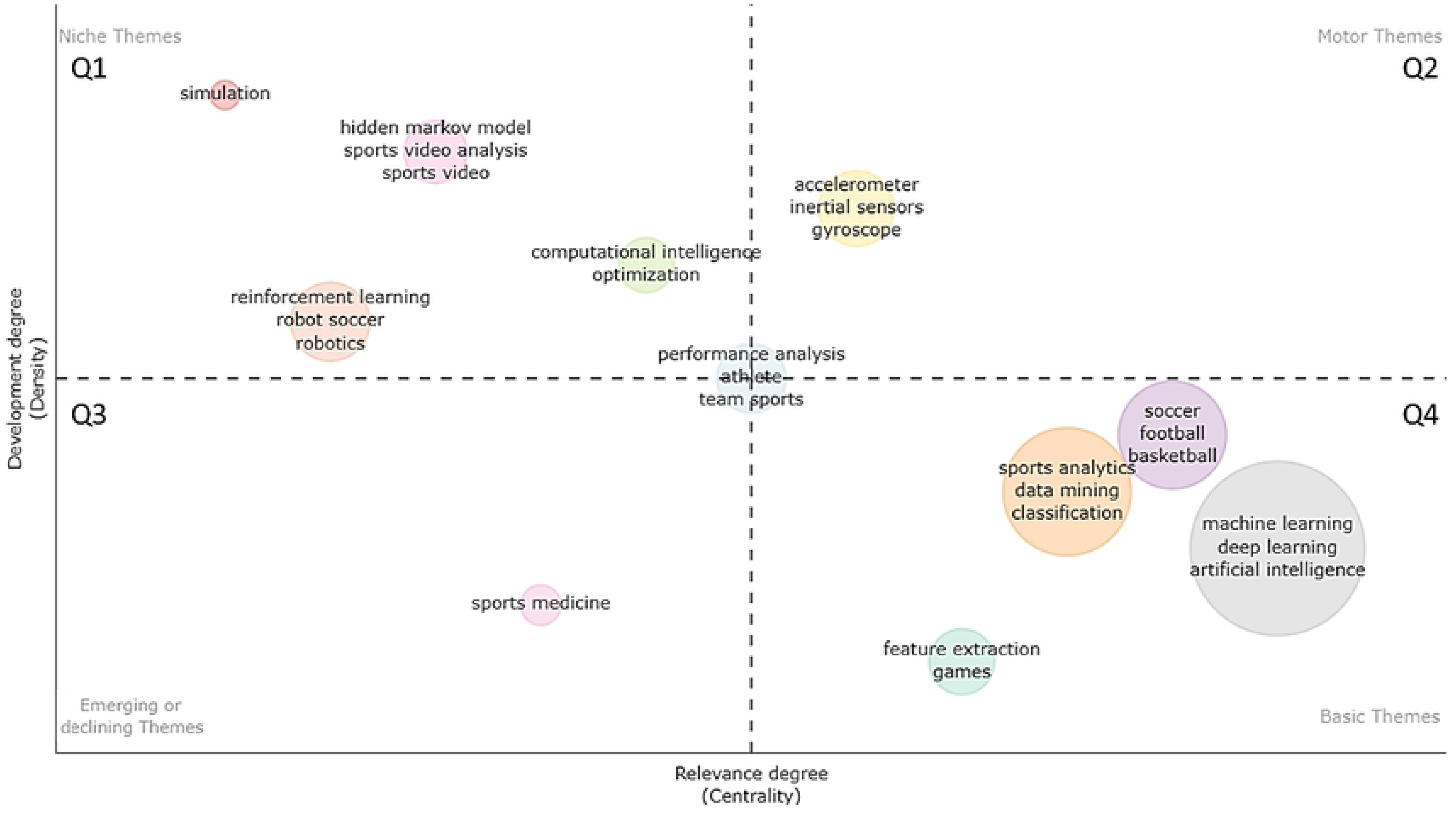
Thematic map using Walktrap for clustering: The size of the spheres maps the total number of documents to each theme. The top three most frequently occurring labels are displayed per cluster. The more relations a node shows with others in the network, the greater its centrality and importance. The density representing the cohesiveness among the nodes gives clues as to the capability of a research field to develop and sustain itself

**Fig 13.**
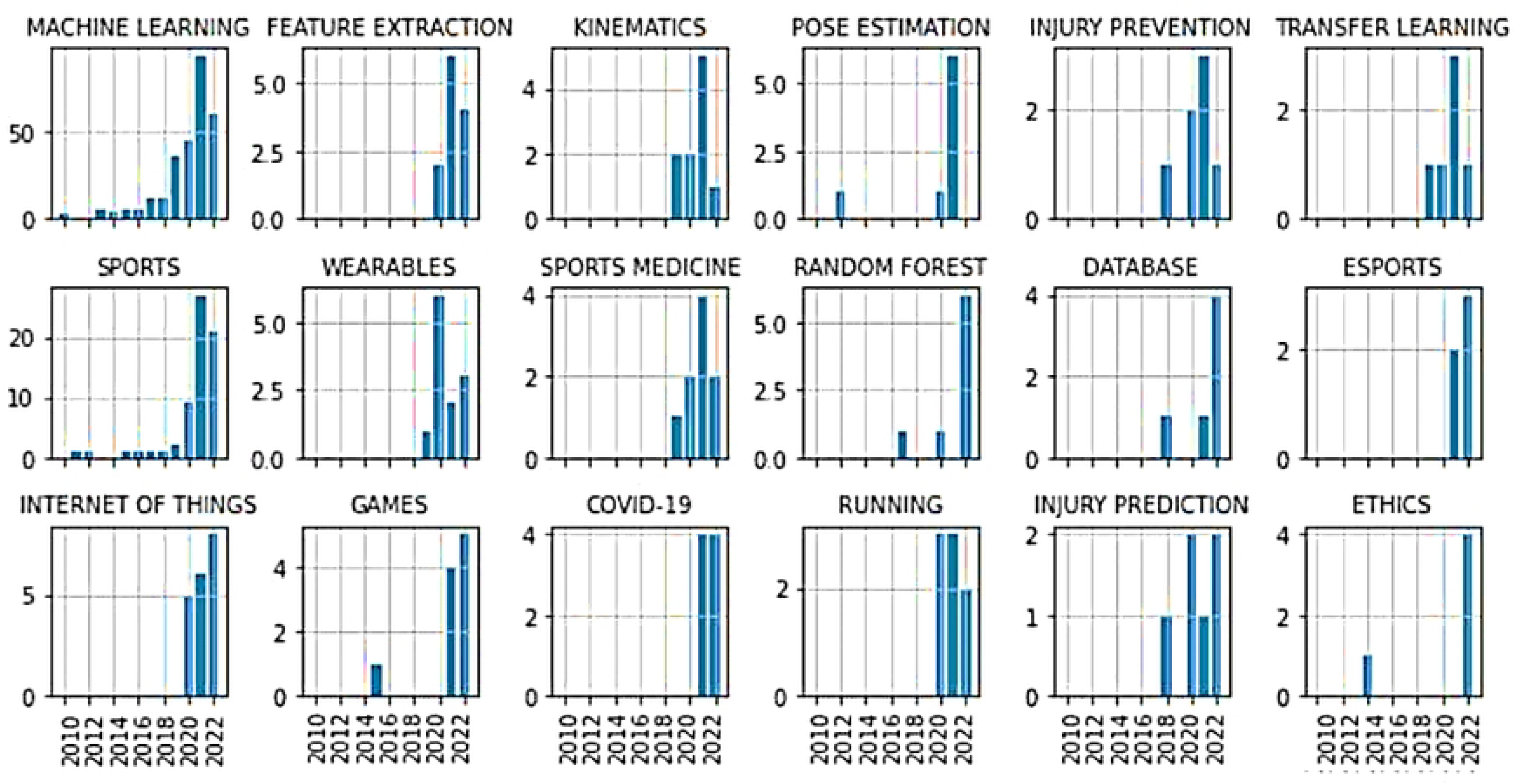
Trend topics using author keywords: Keywords are displayed that obtained more than 80% of their total count in the years 2020–2022. For comparison, the keywords “machine learning” and “sports” are additionally displayed on top. Each y-axis maps the total number of occurrences.

Overall, according to the meaning of the dominant keywords, a high overlap with the contents of the most cited publications listed in Table 1 could be observed. It is notable that when the topics that are covered by the keywords or the most frequently cited publications are assigned to sub-disciplines of sports science, their primary focus is on sports informatics, training science, biomechanics, and sports medicine. Research activities in the fields of sports pedagogy, sports sociology, and sports economics were not evident from our analyses.

Figure 12 presents the corresponding thematic map of AI research in sports. The themes are divided into four quadrants by the horizontal and vertical dashed lines: motor/driving themes (upper right; Q2), basic/underlying themes (lower right; Q4), niche/very specialized themes (upper left; Q1), and disappearing or emerging themes (lower left; Q3). Themes in Q4—such as in general AI (inclusive of its subareas), different specific sports, and subfields of the AI pipeline—are basic, foundational themes that are very important for the field’s general development, characterized by high centrality but low density. Various measuring technologies (e.g., accelerometers, inertial sensors, gyroscopes) can be subsumed under the driving themes of Q2; this makes sense because they often provide complex data, making AI techniques more and more relevant due to static inference methods being limited in those contexts [95]. Looking at Q3, including the subsequent results of trending topics, sports medicine can be identified as an emerging theme. The themes of Q1 (e.g., “simulation”, “hidden Markov model”) are highly specialized and rare, creating internal bonds, as indicated by their low centrality but high density. The theme “performance analysis” is located between all quadrants, indicating that some aspects are basic and necessary for file development, some are emerging or declining, some are fundamental drivers, and some are highly specialized niche themes.

Overall, it is notable that the manual grouping of the most frequently occurring keywords fits with the data-driven clustering of the thematic map to a high degree; this also speaks to the coherence of the obtained solution.

Figure 13 shows the evolution of author keywords, which has attracted particular attention in recent years. In addition to the neural networks that have been in use for a long time, as well as the support vector machine (see Figure 9), the random forest algorithm has recently been given special attention. This is consistent with the results of the co-citation analysis, where a large cluster was found around the work of Breiman (2001) [82]. According to methodological aspects in the context of the AI workflow, attention has also recently been paid to “feature extraction” and “transfer learning”.

The biomechanical aspects of “pose estimation” and “kinematics” have only recently attracted particular attention. Furthermore, the field of “sports medicine” and the matching topic of injury prevention are mentioned frequently—especially recently. The sport of running and e-sports have also received attention in connection with AI in recent years.

Unsurprisingly, “COVID-19” also attracted attention in 2021 and 2022 during the period when the world was still struggling deeply with the pandemic. The future infection situation will probably be decisive for the progression or decline of this topic. Another aspect that has emerged—especially in 2022—is the ethical discussion of AI in sports. Please refer to Figure 11 for information on further trend topics.

At this point, it should be noted that our analysis probably does not depict topics that have arisen very recently. For example, few recent publications have addressed the black-box character of AI models [96, 97], which does not comply with the General Data Protection Regulation (GDPR) [98], using explainable AI. However, this theme does not appear as a trend topic using the author keywords at present, although it may be visible in a bibliometric analysis in the next few years.

## Limitations and further research

In addition to the aforementioned advantages for our research aims, the bibliometric methodology is also associated with some inherent limitations. The analysis is based on technical decisions, e.g., the selection of English as the search language or the focus on the highest-quality papers (i.e., articles and reviews), which may have resulted in the exclusion of other research papers. Therefore, future research should also consider the inclusion of other document types, e.g., conference papers or preprints, as these may provide better insights into the very latest research themes because their publication times are often shorter. Articles in anthologies or monographs could also be interesting, since social science sub-disciplines of sport use them as publication opportunities. Furthermore, it has been shown that different databases (e.g., Google Scholar, Scopus, Web of Science) have certain differences in their search results [99]. Therefore, a multisource comparison and mutual supplementation of the different databases could result in a broader literature overview. Reported issues with the ambiguity of author names could be addressed in the future by using ORCID IDs to prevent potential incorrect merging of authors with the same first and last names [81].

In addition, the used search terms could be expanded to include further aspects that are associated with the term “sports”, in order to identify publications that deal with topics that can be assigned to sports without mentioning the term itself (e.g., physical activity, exercise, training). At this point, the problem of defining the term sports should not go unmentioned. One should be aware that, especially in everyday life, there are a wide variety of (individual and cultural) ideas and definitions of sports—and this applies not only to the term “sports”, but also to terms that are often associated with sports, such as the term “training” [100, 101]. For example, a central point of discussion is whether e-sports constitute sports [102]. This problem was dealt with in this study in such a way that, in simplified terms, it was assumed that publications containing the term “sport*” could be assigned to the research interest of sports science. Therefore, publications from the field of e-sports or—in some cases—from the field of board games (e.g., “Checkers is solved” [74]) were also included. It should be noted that, depending on the individual or context-specific understanding of the definition of sports, these publications might also be classified as unrelated to sports.

## Conclusion

The present study provides a detailed analysis of the leading journals, authors, institutions, keywords, and research themes associated with interconnections between the fields of AI and sports. However, it seems that AI has not yet taken a leading role in sports science but has been gaining more and more importance recently, and sport is an area of application for AI. Our key findings are as follows: (a) The literature and research interest concerning AI and its subcategories is growing exponentially, with considerable potential for further growth in the coming years. (b) The top 20 most cited works account for 32.52% of the total citations, including the presence of “hot papers”, which are still relatively young and have already received considerable numbers of citations. (c) The top 10 journals are responsible for 28.64% of all published documents and indicate a trend of interdisciplinary research. (d) In addition to strong collaborative relationships, delimited small collaboration networks of individual institutions are identifiable. (e) The three most productive countries are China, the USA, and Germany; collaborations appear to be taking place more within the clusters of high- and middle-income countries. (f) Different research themes can be characterized using author keywords, with potential current trend topics including the fields of biomechanics, injury prevention or prediction, new algorithms, and learning approaches. AI research activities in the fields of sports pedagogy, sports sociology, and sports economics seem to have played a subordinate role so far.

These findings provide a better understanding of the research situation regarding the use of AI in sports and may aid researchers in identifying currently relevant research topics, as well as research gaps, through revealing fields of research with much evidence and those that have only been considered marginally thus far. Furthermore, they may be helpful in improving the obviously necessary networking in the context of interdisciplinary and transdisciplinary research, in order to combine competencies from different disciplinary approaches. Hence, this work could be a starting point for systematically initiating stronger networking between authors and their countries/institutions, which might offer yet-untapped potential in their interconnection according to the present study’s results.

Today, AI seems to play an important role not only in sports research, but also in sports practice. Numerous companies (e.g., Apple, Microsoft, Enduco, iDIERS, Aaptiv) are advertising commercial offers that use AI technologies and, thus, have partially opened up areas that research has not yet sufficiently investigated. Therefore, it seems important that research does not lag behind in questioning these economic interests with scientific evidence.

## Footnotes

1 Hereafter, the term artificial intelligence (AI) is used even when individual publications refer specifically to machine learning (ML) or deep learning.

